# Single-particle track colocalization using CoPixie reveals the impact of cancer-associated POT1 mutations on telomerase-telomere interactions

**DOI:** 10.1101/2024.02.19.580537

**Authors:** Samuel Prince, Kamélia Maguemoun, Mouna Ferdebouh, Emmanuelle Querido, Amélie Derumier, Pascal Chartrand

**Author notes:** Present address: Department of Biology, Queens University, Kingston, Ontario, Canada (Kamélia Maguemoun). Present address: de Duve Institute, Université Catholique de Louvain, Brussels, Belgium (Amélie Derumier). The first three authors should be regarded as joint First Authors.

## Abstract

Single-particle imaging and tracking can be combined with colocalization analysis to study the dynamic interactions between macromolecules in living cells. Indeed, single-particle tracking has been extensively used to study protein-DNA interactions and dynamics. Still, identification and quantification of binding events at specific genomic loci remains challenging. Herein we describe CoPixie, a new software that identifies colocalization events between a theoretically unlimited number of imaging channels, including single-particle movies. We employed CoPixie with live cell single-molecule imaging of telomerase and telomeres to test the model that cancer-associated POT1 mutations facilitate telomere accessibility. We show that OB-fold mutants POT1-ΔOB, Y223C, D224N or K90E increase telomere accessibility for telomerase interaction and the cumulative dwell-time of telomerase at telomeres. However, unlike POT1-ΔOB or D224N mutants, the POT1 Y223C and K90E mutations also increase the duration of long-lasting telomerase interactions at telomeres. Our data reveal that telomere elongation in cells expressing cancer-associated POT1 mutants arises from the dual impact of these mutations on telomeres accessibility and telomerase retention at telomeres. CoPixie can be used to explore a variety of questions involving macromolecular interactions in living cells, including between proteins and nucleic acids, from multi-color single-particles tracks.

## INTRODUCTION

Colocalization analysis constitutes a key technique to study and quantify macromolecular interactions in cellular biology. Both pixel-based and object-based colocalization analyses have been developed over the years to provide reproducible and quantitative information on macromolecular interactions within cells (1). Although this technique is limited in its capacity to identify new interactions between macromolecules, it provides information on the dynamics and the cellular context of these interactions. Colocalization analysis thus complements genetic, biochemical and biophysical assays in our understanding of cellular interactions between macromolecules (2).

The advent of single-molecule imaging at nanometer-scale resolution changed our perspective on macromolecules dynamics in living cells (3). While several algorithms have been developed to quantify colocalization between single-molecules in fixed cells (4,5), their application to study colocalization events between molecules in living cells is challenging. Due to the rapid rate of acquisition and the sparse labeling that limit the number of molecules that are tracked in living cells, few colocalization events between proteins or protein-nucleic acids can be detected, and most interactions have to be inferred from changes in the diffusion coefficient of the particles (6) or in their residence time (7).

Colocalization analysis using multi-wavelength single-particle tracks in living cells represents a more advanced alternative to identify specific interactions as it combines both spatial and temporal information. This approach has been mostly used to study the colocalization and stoichiometry of receptors at the surface of plasma membrane (3,8,9) or the binding of transcription factors to chromatin (10). Current strategies to quantify colocalization and binding events using dual-color single-particle tracking in living cells include the identification of correlated tracks between two channels (11,12), the identification of coincident tracks (13), or by performing particle tracking only in the region of interest tagged with a fluorescent-labeled gene array (14). However, for object-based colocalization analysis, colocalization events are often scored manually, which limits the throughput of the analysis (10,15). So far, few tools are available to detect and quantify combined track and object-based colocalization events between two molecules or more in living cells using multi-wavelength single-particle tracking.

The repetitive nature of telomeric sequences and their association with a specific complex of proteins named Shelterin ensure that telomeres can be easily labeled with fluorescent proteins and imaged as single foci by live-cell microscopy (16,17). Telomeric repeats are replenished by a specialized enzyme called telomerase, a ribonucleoprotein complex with reverse transcriptase activity (18,19). In recent years, single-molecule imaging has been applied to study telomerase dynamics and binding to telomeres in living cells. Cell lines expressing the reverse transcriptase hTERT endogenously tagged with Halo-tag (20,21) or MS2-tagged hTR telomerase RNA (22), have been used to image single molecules of Halo-TERT or MS2-hTR RNA in living cells. This opened the possibility to use multi-wavelength single-molecule imaging to study the regulation of telomerase-telomere interactions *in vivo*. Indeed, single-molecule imaging was used to explore the role of the telomeric single-strand DNA-binding protein POT1 in the negative regulation of telomerase binding at telomeres (22).

Heterozygous dominant mutations in POT1 have been identified in several cancers. These mutations lead to chromosomal abnormalities, such as telomere fragility and end-to-end chromosome fusions (23). However, the ability of cancer-associated POT1 mutations to promote telomerase-dependent telomere elongation was suggested to be the prime driver of cancer progression (24,25). A current model posits that these mutations disrupt POT1-driven inhibition of telomerase access to telomeres, although this model has not yet been validated *in vivo* (26,27). Single-molecule imaging of telomerase at telomeres may help resolve this question, but the lack of robust algorithms to analyze multiple colocalization events between telomerase particles and telomeres limits the throughput of such studies.

Herein, we introduce CoPixie, a program that identify co-localization events between a theoretically unlimited number of imaging channels, including from multi-wavelength single-particle tracking. Since CoPixie also provides temporal information on the colocalization events, the interaction dynamics between two molecules can be obtained *in vivo*. We demonstrate its capabilities by studying the impact of cancer-associated POT1 mutations on the dynamic interaction between telomerase and telomeres using dual-color single-molecule imaging and particle tracking in living cells.

## MATERIAL AND METHODS

### Cell culture procedures and treatments

HeLa hTR^5’MS2^ cells are described in (22) and were cloned from parental HeLa 1.3 cells with long telomeres, as identified by the de Lange lab (28). These cells express mCherry-TRF1 to detect telomere foci, and mCherry-CDT1 to mark cells in S-phase or G2 for both filming and particle tracking in the CoPixie analysis pipeline. Cells were cultured in Dulbecco’s Modified Eagle Medium (DMEM, Corning) supplemented with 10% Fetal Bovine Serum, 2 mM L-glutamine, and 100 U/ml Penicillin-Streptomycin. Transduction of myc-tagged POT1 mutants was achieved using lentiviral vector infection, followed by puromycin selection for 3 days. Cell lines stably expressing POT1 were maintained for no more than five weeks after selection.

### Western Blot analysis

Cell pellets were lysed in a standard RIPA buffer with sonication, as described previously (22). Protein concentration was quantified with the BCA Protein Assay Kit (ThermoFisher Pierce) and 20µg of protein/sample was loaded per lane on SDS-PAGE gel. Primary antibodies used were anti-c-myc (11667149001, 9E10 mAb from Roche; 1:250), anti-α-Tubulin (T5168 Sigma; 1:5000) or anti-β-Tubulin (480011 Invitrogen;1:10,000). Secondary antibodies used were either anti-rabbit (170-6515 Bio-Rad; 1:3000) or ECL anti-mouse (NA931 Cytiva; 1:5000).

### Immunofluorescence

POT1-myc and mCherry-TRF1 expressing cells were plated on 22×22 mm glass coverslips and fixed with 4% formaldehyde. The cells were permeabilized in 1X PBS with 0.5% Triton X-100, then blocked in 1X PBS with 0.1% Triton X-100 and 2% BSA at room temperature. Coverslips were then incubated with the anti-myc antibody (11667149001 9E10 mAb from Roche; 1:250 in blocking buffer) for two hours at room temperature. After washes, coverslips were incubated with a Alexa Fluor 488 anti-mouse antibody (715-585-150 from Jackson Immuno Research 1:7000). Coverslips were mounted with Vectashield (Vector Laboratories) containing DAPI for nuclei staining. Z-stacks were acquired with a 100X NA 1.3 oil objective on a Zeiss Axio-Imager Z2 upright widefield epifluorescence microscope equipped with a sCMOS camera (2048×2028 pixels). Images were deconvoluted with the Zen 2.6 Pro software (Zeiss).

For immunofluorescence on the POT1 Y223C and D224N mutants, a secondary Alexa Fluor CF680 anti-mouse antibody (A-21057 Invitrogen 1:1000) was used instead. Z-stacks were acquired with a 100X NA 1.46 oil objective on a Zeiss Axio-Observer Z1 Yokogawa CSU-X1 spinning disk inverted confocal microscope equipped with two EMCCD Evolve cameras (Photometrics, 512×512 pixels). Maximal intensity projections of nuclear Z-slices were generated in ImageJ for illustrative purposes for all the POT1 immunofluorescence images.

### Live-cell imaging of hTR and telomeres

For live-cell microscopy, cells were grown on glass-bottom 35mm dishes (Fluorodish FD35-100 World Precision Instruments) in growth media. Three hours before imaging, growth media was replaced with imaging media (DMEM phenol red-free with 10%FBS and 25mM HEPES pH7.4). Cells were filmed on a Zeiss Axio-Observer Z1 Yokogawa CSU-X1 spinning disk inverted confocal microscope equipped with two EMCCD Evolve cameras (Photometrics, 512×512 pixels, 16 µm). The images were acquired with a 100X NA 1.46 oil objective. The pixel size of the images obtained was 0.133µm. The Zeiss TempModule was set to 37°C two hours prior to imaging in order to allow all components of the imaging chamber to reach temperature. The two lasers, a 488nm 100mW diode, and a 561nm 40mW diode, were turned on for a minimum of one hour before imaging. For dual-camera imaging with the 488nm and 561nm lasers, emission was split by a Zeiss FT 560 beam splitter. The Zeiss DualCamera Wizard alignment software was calibrated with a slide of multi-fluorescent tissue to apply an Affine (translation + rotation + isoscaling) transform to the 561nm laser image. The czi files were exported to the OME TIFF format from within the ZEN software to preserve the Affine alignment of the second camera images in ImageJ.

For short hTR-telomere interaction movies, 100 images were acquired continuously in dual-camera mode with a 70ms exposure time (plus 30 ms transfer, resulting in a 100 ms interval). For long hTR-telomere interaction movies, mCherry-TRF1 and hTR-GFP were acquired sequentially on a single Evolve camera at 2 second intervals for 150 images (300 seconds). GFP was acquired first with 100ms exposure time and mCherry with 150ms exposure time. Focus was maintained with the Zeiss Definite Focus LED IR 835nm laser every 10 images. The hTR-telomere movies were analysed with ImageJ TrackMate using a custom macro (available in Supplementary Methods) that includes an exponential fit bleach correction applied to each image series. Briefly, the TrackMate LoG spot detection was set to a diameter of four pixels (0.53 µm)for both telomeres and hTR particles. The simple LAP tracker was used with a linking max distance of 0.8 µm, and zero gap-closing for hTR. For telomeres, linking max distance was 0.2 µm and the max frame gap was 3. Tracks comprising of 5 or more spots were kept in the TrackMate output.

### CoPixie analysis of hTR-telomere interactions

CoPixie can be installed from https://github.com/drs/CoPixie. The hTR-telomere particle tracking data files were analysed with CoPixie for colocalization events as detailed in the Supplementary Methods. hTR particles were defined by a centroid and a square of 3×3 pixels (size 400×400nm) which covers the same area as the hTR particles in the image series. Telomeres were defined by a centroid and a mask to map the actual size of each telomere. Data aggregation was performed within CoPixie with a custom Python script written for hTR-telomere interaction dynamics (file available upon request). The aggregation script applied a criteria that the telomere tracks persist for 50% or more of the movie frames, to avoid counting the same telomeres twice. A second criteria was applied so that only hTR-telomere interactions lasting 5 frames or more for the short (10 Hz) movies (>0.5 seconds) and 5 frames or more for the long (0.5 Hz) movies (>10 seconds) were selected. The aggregation script calculated the last frame of interaction minus the first frame between each pair, which resulted in the dwell-time of the hTR particle at the telomere. The script also calculated the cumulative dwell-time of hTR particles for each telomere.

### Dual-camera imaging and tracking of fluorescent beads for CoPixie validation

TetraSpeck microspheres (ThermoFisher T7279 0.1µm) suspended in 0.1% gelatin were plated on a glass-bottom Fluorodish. The fluorescent beads were excited simultaneously with the 488nm and 561nm lasers, then emission was split with the FT 560 beam splitter and sent to the two Evolve cameras as described in the previous section. Images were acquired continuously with 30ms exposure time (plus 30 ms transfer, resulting in 60 ms interval) for 100 images. An Affine transform was applied to the 561nm image to align it to the 488nm image. The localization precision on TetraSpeck microspheres after this Affine correction was previously measured to be 45 nm (22). The ImageJ TrackMate plugin was used for single particle tracking of each channel with the LoG spot detection set to four pixels (0.53 µm). The simple LAP tracker was used with a linking max distance of 0.8 µm, and zero gap-closing. Tracks comprising of 7 or more spots were kept in the TrackMate output.

### Mathematical and Statistical analyses

GraphPad Prism was used for survival probability analysis (1-CDF) of hTR dwell-time at telomeres from single-particle tracking experiments performed at 10 Hz or 0.5 Hz. Curve fitting was performed with an equation for two-phase exponential decay using least squares regression and weighting by 1/Y^2^ for optimal curve fitting (R^2^ > 0.99). The following constrains were imposed on the parameters: plateau was fixed at 0 and KSlow > 0. The residence time (τ_fast_ and τ_slow_) and half-life (fast and slow) were derived from the model. Values are reported as mean ±95% Confidence Interval (CI). Comparison and statistical analysis of residence time and half-life of hTR particles at telomeres from the various cell lines was performed with multiple unpaired Student t-test. For the cumulative dwell-time analysis, data were fitted to a Kaplan-Meier survival curve using GraphPad Prism. Comparison and statistical analysis of survival curves was performed using a Log-rank (Mantel-Cox) test.

## RESULTS

### Description of CoPixie

CoPixie is a Python script that identifies colocalization events between single-particle tracks in multiple channels (Figure 1). The program is available for Linux, Windows and MacOS (https://github.com/drs/copixie). CoPixie handles image processing in batch for multiple datasets, according to a configuration file provided by the user, as detailed in the Supplementary Methods. Particle position information is generated for each channel separately using single-particle tracking (for diffraction-limited particles) and masks (for larger particles or cellular compartments), then provided to CoPixie for colocalization analysis. CoPixie was designed to work with 2D particle tracking files generated by the ImageJ TrackMate plugin (29), but it could be used with any files containing particle centroid coordinates.

**Figure 1:**
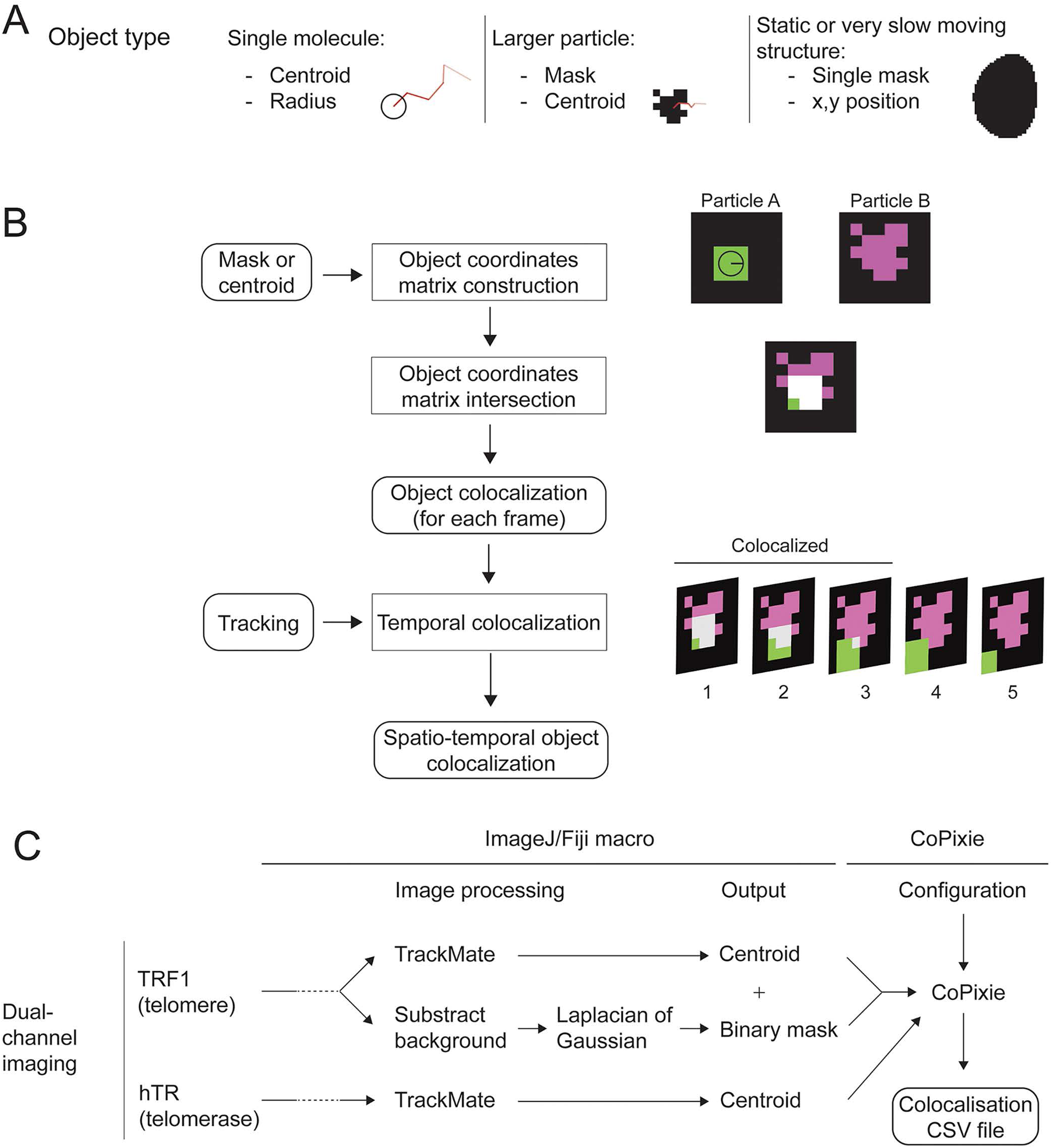
Description of CoPixie and the hTR-telomerase interaction pipeline. **A**) The object types used by CoPixie; a) a centroid plus radius for single molecules, b) a mask plus a centroid for larger particles, and c) a single mask plus an x,y position for slow moving structures. **B)** Schema summarizing the sequential steps of the CoPixie script. **C)** Description of the pipeline for the analysis of colocalization events between hTR and telomeres using ImageJ and CoPixie.

The CoPixie object types are described in Figure 1A; 1) centroid and radius, 2) mask and centroid, or 3) static mask plus x,y position. Indeed, an innovative CoPixie feature is the ability to use a still image for one of the imaging channels. The architecture of the CoPixie matrixes allows the program to identify colocalizations events between a theoretically unlimited number of imaging channels. CoPixie scores colocalization based on pixel overlap, which is well adapted for single molecule imaging using pixel-based digital camera images. In many single-molecule imaging applications, particle area is diffraction-limited and signal-to-noise ratios are low, thus colocalization analysis using exclusively mask-based algorithms would be challenging. Colocalization criteria which use centroid-to-centroid distance of the two objects may not give precise results for larger particles with irregular shapes. Since CoPixie allows both types of objects, it was designed to offer both flexibility and robustness in colocalization mapping.

In CoPixie, single molecules objects are defined by a radius around the centroid which defines a virtual square, usually set to 3×3 pixels, with pixel dimensions matching the image pixels (see Supplementary Methods for details). For larger particles with an irregular shape, the objects are defined both by a mask derived from image segmentation and by the centroid obtained for particle tracking. CoPixie scores a colocalisation event in the image frame when at least one pixel of the two objects overlaps, as illustrated in Figure 1B. The script consists of two main steps: (1) object colocalizations are identified frame-by-frame using the intersection of the coordinates of every object, and (2) particle tracking information is used to link colocalizing objects in time, which allows the user to extract metrics such as the length of the interaction. The output of CoPixie consists of tables of colocalization events for each dataset. Notably, CoPixie identifies which segments of particle tracks colocalize and ouputs information on any gaps in colocalization.

### Validation of CoPixie

To demonstrate the capacity of CoPixie to perform colocalization of single-particle tracks, we first used multicolor fluorescent beads since they should yield perfect colocalization between channels. The 0.1 micron beads diffusing in a viscous solution were excited with the 488 nm and 561 nm lasers for simultaneous image acquisition with two EM-CCD cameras. Tracks were generated indepently for each channel with ImageJ TrackMate. As expected, overlay of the 488 nm and 561 nm channels revealed overlapping tracks coming from single beads (Figure 2A _Matching). Although each bead was labeled with both fluorophores, some tracks or track segments were obtained in one channel but not in the other, possibly because of lower signal-to-noise ratio images in one channel. CoPixie colocalization analysis of 324 tracks in the 488nm channel and 369 tracks in the 561nm channel captured in eight different movies revealed that a median of 87% of tracks were colocalized with another track in the other channel (Figure 2B_Matching). All the missing colocalizations came from tracks that did not have a corresponding track in the other channel (see Supplementary Methods for full details of the CoPixie settings). To assess the level of random track colocalization, images from non-matching movies were taken (*i.e* the 488 nm channel of one movie was paired to the 561 nm channel from another movie) and analyzed with CoPixie. As expected, few colocalization events between tracks were detected, and they lasted for only 1 or 2 timepoints (Figure 2A and B non-matching).

**Figure 2:**
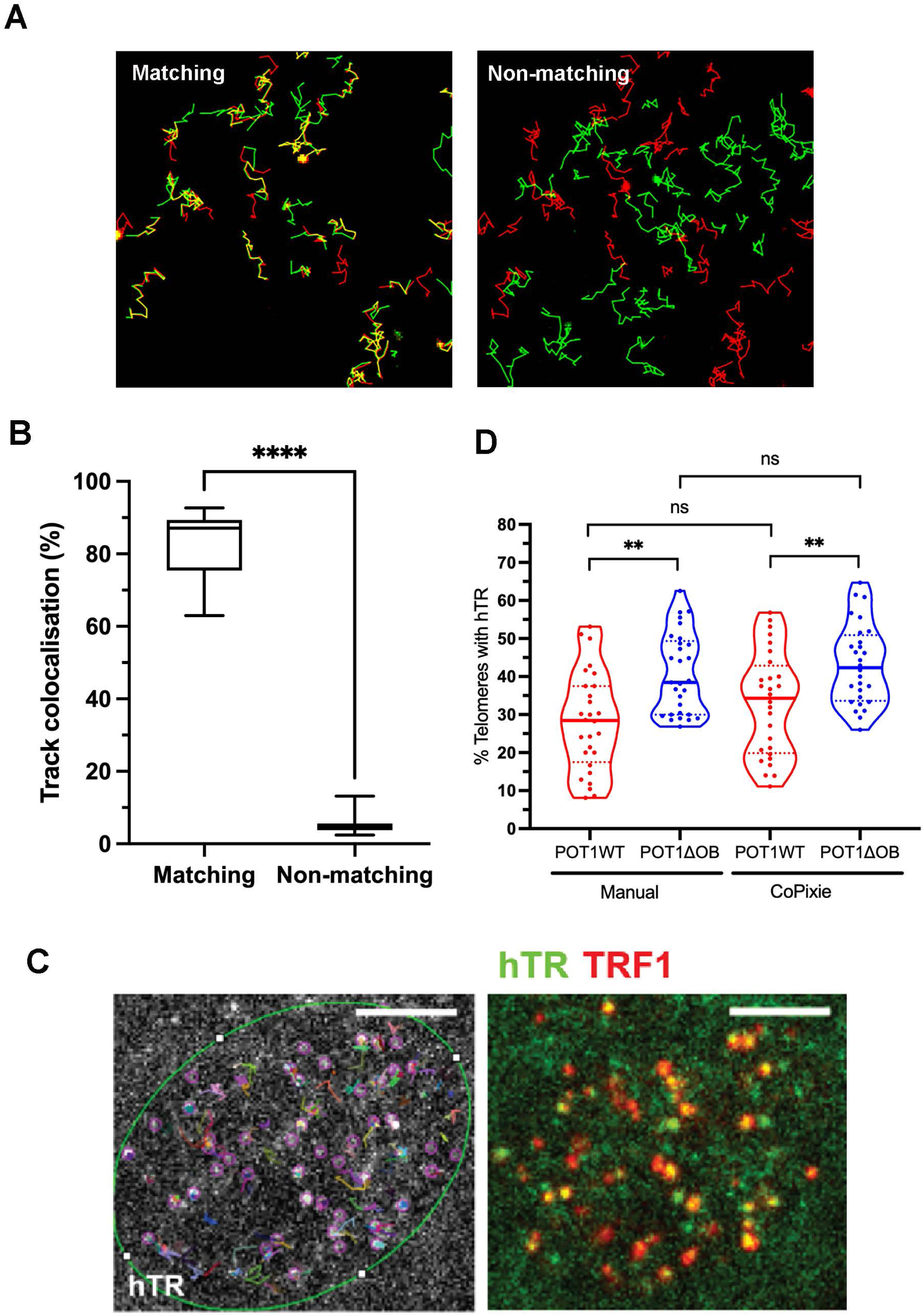
Validation of CoPixie. **A**) Tracks from fluorescent beads imaged by dual cameras and analyzed with TrackMate. Overlay of tracks from the two channels of the same movie, matching (left), and tracks from different movies, non-matching (right). **B)** Percentage of track colocalizations for matching and non-matching movies as measured by CoPixie. ****p<0.001. **C)** Particles of hTR are detected (violet circles) and tracked using TrackMate. Overlay of a single image of hTR (MCP-sfGFP, green) and telomeres (mCherry-TRF1, red). Scale bars: 5 μm. **D)** Data from Laprade et al. (2020) were re-analyzed using the CoPixie pipeline. N= 29 cells expressing POT1-ΔOB and 28 cells expressing POT1-WT. **p < 0.01; ns: not significant.

CoPixie can also quantify track colocalization events that are located in a specific region of an image series. Supplementary Figure 1 shows the three regions of a circle tested for this demonstration: the border, the center, and outside of the circle. CoPixie was able to accurately identify the track segments which colocalized both with each other and with a specific region. This demonstration highlights CoPixie’s potential use to monitor single-molecule localization in sub-cellular organelles or membranes.

To further validate CoPixie, we determined whether the program could reproduce colocalization results from a published dataset of colocalization events between single particles of the telomerase RNA hTR and telomeres, previously quantified by manual analysis (22). To image single particles of hTR, HeLa 1.3 cells with three MS2 stem-loops integrated at the 5’ of hTR (HeLa hTR^5’MS2^) and stably expressing the MS2 coat-protein fused to superfolder GFP (MCP-sfGFP) and mCherry-TRF1, enable the detection of both telomeres and hTR particles using a spinning disk confocal microscope (Figure 2C). Imaging was performed to compare hTR-telomere interactions in cells overexpressing hTERT, and POT1-WT or a N-terminal deletion of the first OB-fold domain of POT1 (POT1-ΔOB). In our previous manual analysis, colocalization events between hTR and telomeres were scored using the TrackScheme view of hTR track positions during TrackMate particle tracking. However, this manual scoring of hTR-telomere interactions was a lenghty process, averaging 25 minutes per nucleus. Our published results using manual analysis revealed that a median of 28% of the telomeres were visited by hTR in cells expressing POT1-WT and 41% in cells expressing POT1-ΔOB, *i.e*. an increase of 13 ± 3% (difference between means ± SEM) of hTR colocalizations at telomeres in the presence of the POT1-ΔOB mutant (Figure 2D). With the same dataset, a similar distribution was obtained with CoPixie, which calculated that a median of 33% of the telomeres per cell were visited by hTR in cells expressing POT1-WT compared to 43% in cells expressing POT1-ΔOB; a difference of 10 ± 3% (difference between means ± SEM) between the two cell lines (Figure 2D). Considering the standard error of the mean (SEM) of 3%, the difference between the means obtained by manual colocalization (13%) versus CoPixie (10%) are within the confidence intervals. Differences in the results from POT1-WT (or POT1-ΔOB) between CoPixie and manual colocalization were statistically non-significant. Furthermore, the same upward shift in the percentage of telomeres colocalized with hTR was observed in cells expressing POT1-ΔOB compared to cells expressing POT1-WT in both analyses (Figure 2D). Altogether, these results show that CoPixie can accurately detect and quantify colocalization events from dual-color single-particle tracking data.

### Application of CoPixie to telomerase particle tracking and colocalization with telomeres in the context of cancer-associated POT1 mutations

It is well established that the shelterin component POT1, which binds single-stranded telomeric repeats, acts as a negative regulator of telomerase activity at telomeres (30,31). However, *in vitro* evidence show that TPP1-POT1 can also act as an activator of telomerase processivity (32). Expression of dominant POT1 mutants, including those frequently found in human cancers, leads to rapid telomere elongation in cancer cells (23). Although the current model suggests that most cancer-associated mutations in POT1 fail to inhibit telomerase access at telomeres (23,27), no studies have experimentally validated this model *in vivo*. So far, colocalization studies using RNA FISH showed no impact of cancer-associated mutations in POT1 on hTR recruitment to telomeres (26). Since our previous work using single-molecule imaging of hTR-MS2 revealed that deletion of the first OB-fold domain of POT1 (POT1-ΔOB) increases telomerase access to telomeres (22), we sought to apply high-troughput CoPixie colocalization analysis to explore the impact of cancer-associated mutations of POT1 on their capacity to inhibit telomerase interaction with telomeres *in vivo*.

While several POT1 mutants have a robust impact on its interaction with telomeric ssDNA *in vitro* (33,34), other cancer-associated POT1 mutants have less obvious effect on the protection of telomeric ssDNA. One such mutant is POT1 K90E, which was identified in patients with cutaneous T cell lymphoma (35). This mutation in the first OB-fold domain of POT1 leads to telomeres elongation and telomere fragility, even though the K90E mutant still binds telomeric ssDNA with similar affinity to wild-type POT1 (35).

To image single particles of hTR, HeLa 1.3 cells with three MS2 stem-loops integrated at the 5’ of hTR (HeLa hTR^5’MS2^) overexpressing hTERT were transduced with lentiviral particles expressing myc-tagged versions of POT1-K90E, POT1-WT, POT1-ΔOB or the empty vector. Expression of the Myc-tagged proteins was confirmed by Western blot (Figure 3A) and their localization at telomeres was validated by immunofluorescence (Figure 3B). Simultaneous imaging of single-particles of hTR and single telomeres was performed by constant acquisition at 100 msec/frame for a total of 10 sec (See Supplementary movies 1 to 4). Individual hTR particles were tracked using TrackMate, and their colocalization with telomeres was quantified using CoPixie. Figure 1C schematizes the CoPixie pipeline used for hTR-telomere interactions which combines centroid-based objects for hTR single particles and mask-based mapping for larger irregularly shaped telomeres. This analysis revealed that the percentage of telomeres with at least one colocalizing hTR particle track in cells overexpressing wild-type POT1 was significantly higher than in cells with empty vector alone (medians of 38% and 33%, respectively) (Figure 3C). As previously reported, expression of POT1-ΔOB resulted in an increased percentage of hTR-bound telomeres compared to POT1-WT (Figure 3C). Interestingly, expression of POT1-K90E also increased the percentage of telomeres associated with hTR particles (Figure 3C), suggesting that this mutant increases the accessibility of telomeres for telomerase binding.

**Figure 3:**
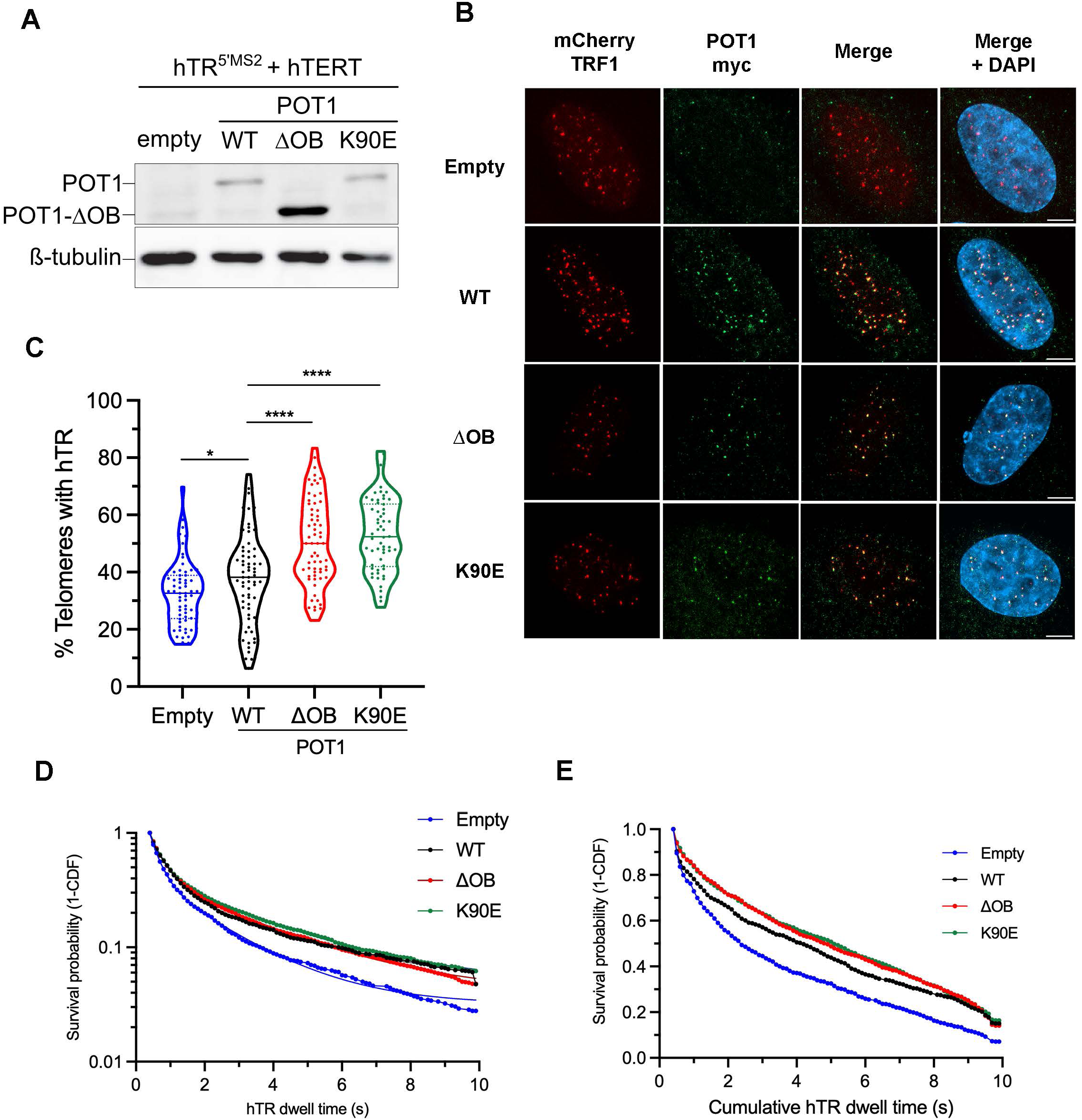
The K90E mutation in POT1 increases telomerase access to telomeres. **A**) Western blot analysis to assess the expression level of myc-tagged POT1 wild-type (WT), POT1-ΔOB, POT1 K90E in HeLa hTR^5’MS2^ cells. β-Tubulin was used as loading control. Empty: cells infected with empty vector. **B)** Immunofluorescence on myc-tagged POT1 WT, POT1-ΔOB and POT1 K90E in HeLa hTR^5’MS2^ cells. Immunostaining was performed with anti-myc antibody (green) and telomeres were detected with mCherry-TRF1 (red). Empty: cells infected with empty vector. DAPI: nuclear DNA. Scale bars: 5 μm. **C)** Percentage of telomeres colocalized with hTR in hTR^5’MS2^ + hTERT cells expressing empty vector, POT1-WT, POT1-ΔOB or POT1 K90E. *p<0.05; ****p < 0.001. N = 58-65 cells. **D)** Survival probability analysis of individual hTR particles at telomeres in hTR^5’MS2^ + hTERT cells expressing the empty vector, POT1-WT, POT1-ΔOB or POT1 K90E. Each curve represents data merged from >1600 tracks from 58-65 cells in 3-4 independent replicates. **E)** Survival probability analysis of the cumulative dwell-time of hTR particles at individual telomeres in hTR^5’MS2^ + hTERT cells expressing the empty vector, POT1-WT, POT1-ΔOB or POT1 K90E. Each curve represents data merged from 700 to 1020 tracks from 58-65 cells in 3-4 independent replicates. Statistical difference between groups was determined using Log-rank test: empty vs WT: p<0.0001; WT vs ΔOB: p=0.005; WT vs K90E: p=0.0023.

The CoPixie data aggregation output also includes information on the dwell-time or residence time of each hTR particle that visits a telomere. Additonnally, the data aggregation script compiled the cumulative dwell-time of all the hTR particles that visited a telomere (*i.e* the sum of the dwell-times of all the hTR particles that visit a single telomere). Fitting hTR dwell-times at telomeres in a survival probability plot using two-phase exponentional decay revealed the residence time and half-life of hTR particles at telomeres in cells expressing POT1-WT, POT1-ΔOB or POT1-K90E, or with the vector control. As shown in Figure 3D, no significant difference in the dwell-time of hTR molecules at telomeres was observed between the various POT1 mutants and wild-type. However, a clear increase in residence time was observed in all the cells overexpressing POT1, either WT or mutants, compared to the control cells infected with the empty vector (Figure 3D and Table S1), showing that POT1 overexpression increases the residence time of telomerase at telomeres, independently of the capacity of POT1 to bind telomeric ssDNA.

The cumulative dwell time of hTR at telomeres, as expected, also showed a significant increase in cells expressing POT1-WT, POT1-K90E or POT1-ΔOB, compared to empty vector. Indeed, the median cumulative occupancy of telomeres increased from 2.4 sec in cells with empty vector to 4.2 sec in cells overexpressing POT1-WT (Figure 3E). Median cumulative occupancy further increased to 4.9 sec in the presence of POT1-ΔOB and to 5.1 sec with POT1-K90E, compared to 4.2 sec for POT1-WT (Figure 3E). Altogether, these results suggest that the dwell-time of hTR molecules at telomeres is not changed in cells expressing POT1-ΔOB and POT1-K90E compared to POT1-WT. However, since the telomeric ssDNA is more accessible in these mutants, more hTR molecules bind to telomeres, which increases the cumulative dwell-time of hTR molecules at each telomere.

### Dual effect of the POT1 K90E mutation on telomerase access to telomeres and long-lasting interactions between telomerase and telomeres

Among the hTR molecules tracked at telomeres, some remained colocalized during the entire 10 second movies, suggesting that they could interact with telomeres for much longer times. To quantify stable, long-lasting interactions between hTR particles and telomeres, we switched to time-lapse acquisition with time intervals of 2 seconds between timepoints for a total of 300 seconds. CoPixie was used to quantify hTR-telomere colocalization events and dwell-time in cells expressing POT1-WT, POT1-ΔOB, POT1-K90E or the empty vector. This imaging mode confirmed that the percentage of telomeres which colocalized with hTR particles increased in cells expressing POT1-ΔOB or POT1-K90E compared to cells expressing POT1 WT (Figure 4A). While a median of 42% of telomeres colocalized with hTR in cells with empty vector or expressing POT1-WT, this percentage increased to 62% in POT1-ΔOB and 59% in POT1-K90E. These results underscore the impact of POT1-ΔOB and POT1-K90E mutations in modulating the accessibility of telomeres for telomerase binding.

**Figure 4:**
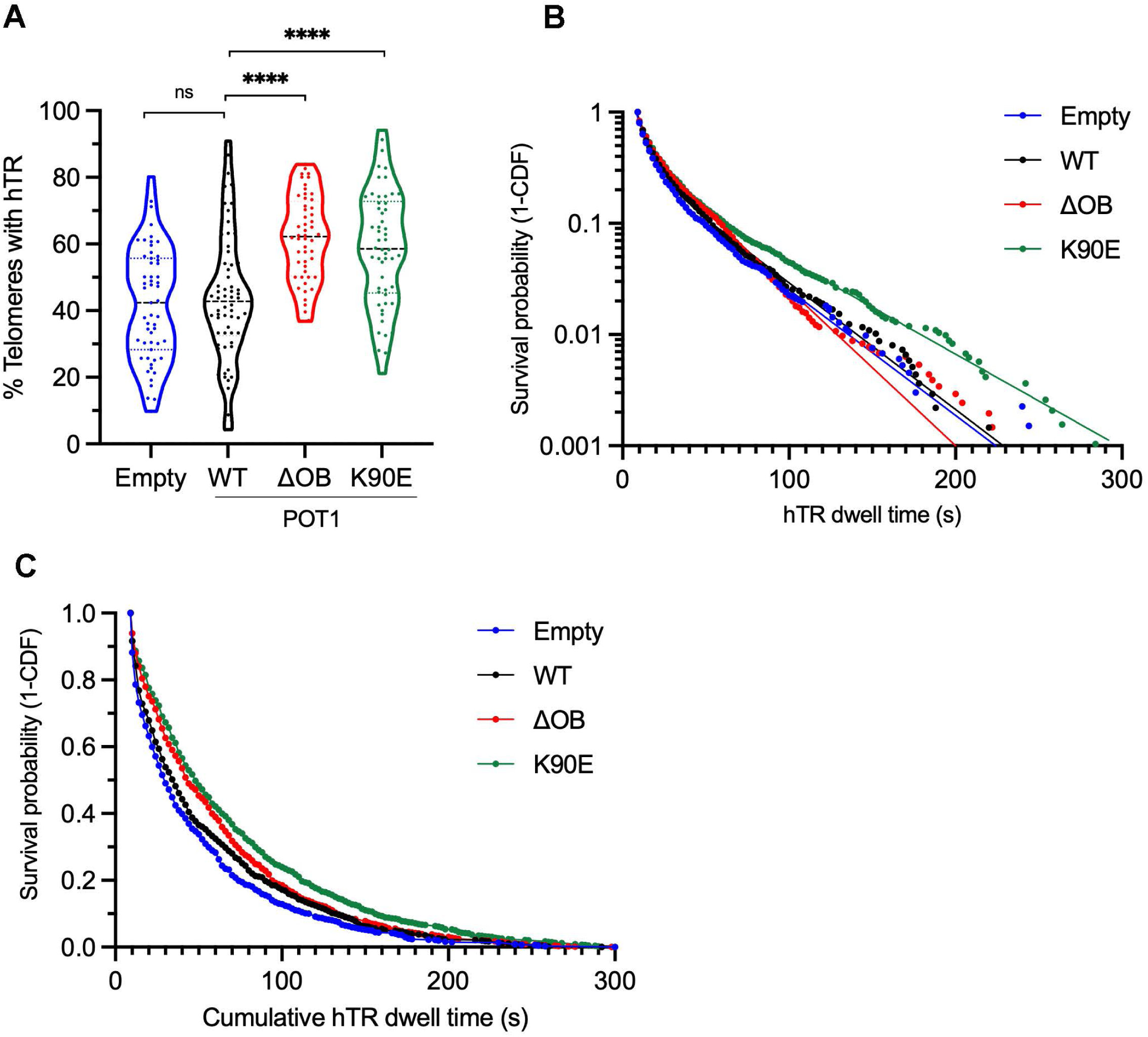
Expression of POT1 K90E increases the long-lasting interactions of hTR with telomeres. **A**) Percentage of telomeres colocalized with hTR particles in hTR^5’MS2^ + hTERT cells expressing the empty vector, POT1-WT, POT1-ΔOB or POT1 K90E. ****p < 0.001; ns: not significant. N = 52 to 62 cells. **B)** Survival probability analysis of individual hTR particles at telomeres in hTR^5’MS2^ + hTERT cells expressing the empty vector, POT1-WT, POT1-ΔOB or POT1 K90E. Each curve represents data merged from 1325 to 2051 tracks from 52-62 cells in 4 independent replicates. **C)** Survival probability analysis of the cumulative dwell-time of hTR particles at individual telomeres in hTR^5’MS2^ + hTERT cells expressing the empty vector, POT1-WT, POT1-ΔOB or POT1 K90E. Each curve represents data merged from 700 to 900 tracks from 52-62 cells in 4 independent replicates. Statistical difference between groups was determined using Log-rank test: empty vs WT: p=0.0268; WT vs ΔOB: p=0.0199; WT vs K90E: p<0.0001.

The dynamics of hTR long-lasting interactions with telomeres was also measured in the 300 second movies. A survival probability plot was generated from hTR binding events to telomeres from cell lines expressing POT1-WT, POT1-ΔOB, POT1-K90E or the empty vector. Survival probability plots were fitted with a two-phase exponential decay model, and binding constants were obtained (Figure 4B and Table S2). No significant difference in hTR long interactions at telomeres were observed between cells expressing POT1-WT, POT1-ΔOB or the empty vector. Surprisingly, unlike POT1-ΔOB, expression of POT1-K90E increased the retention of long-lasting bound molecules of telomerase at telomeres, with residence times increasing from 38,1 to 51,6 seconds (Figure 4B and Table S2). The cumulative dwell-time of long-lasting hTR particles at single telomeres was also measured in cells expressing POT1-WT, POT1-K90E, POT1-ΔOB or the empty vector. The median cumulative occupancy of telomeres increased from 30 sec in cells with empty vector to 36 sec in cells overexpressing POT1-WT (Figure 4C). Median cumulative occupancy was further increased to 44 sec in the presence of POT1-ΔOB and to 48 sec with POT1-K90E, showing that hTR molecules occupy telomeres for longer times in these mutants. Altogether, these results show that the POT1-K90E and POT1-ΔOB mutations increase the accessibility of telomeres for long-lived interactions with telomerase. However, unlike the POT1-ΔOB mutant, the POT1-K90E mutant also increased the residence time of long-lived hTR molecules at telomeres, suggesting a specific property of this cancer-associated mutant in modifying hTR dynamics at telomeres.

### Dual effect of the POT1 Y223C mutation on telomerase access and retention at telomeres

These results raise questions regarding this unique property of the K90E mutation, which surprisingly was not observed in the OB-fold domain mutant POT1-ΔOB, where K90 is located. To further validate this observation, two other cancer-associated POT1 mutants, Y223C and D224N, were tested for their capacity to impact telomerase access and dwell-time at telomeres. The POT1 Y223C mutation was identified in patients with chronic lymphocytic leukemia (33). Expression of POT1 Y223C leads to telomere elongation and fragility. *In vitro*, the Y223C mutant strongly inhibits telomerase activity, but *in vivo*, expression of this mutant leads to telomere elongation (26). Using RNA FISH to assess hTR colocalization with telomeres revealed no difference in its colocalization in POT1 Y223C cells compared to cells expressing wild-type POT1, which suggested that Y223C did not affect the recruitment of telomerase at telomeres (26). The POT1 D224N mutation has been identified in patients with Hodgkin lymphoma or melanoma (34,36). This mutation in the second OB-fold domain reduces the interaction between POT1 and telomeric ssDNA, resulting in telomere elongation and increased telomere fragility (34). However, the impact of this mutation on telomerase activity at telomeres remains unknown. Both Y223C and D224N mutations are heterozygous, suggesting that these mutants act as dominant negatives (33,34). Although both Y223C and D224N disrupt POT1 binding to telomeric ssDNA *in vitro*, their impact on telomerase association with telomeres *in vivo* remains either inconclusive or unknown.

HeLa hTR^5’MS2^ cells overexpressing hTERT were infected with lentivirus expressing myc-tagged POT1-WT, POT1-ΔOB, POT1-Y223C or POT1-D224N. Expression of the myc-tagged POT1 proteins was validated by Western blot (Figure 5A) and their localization at telomeres was confirmed by immunofluorescence (Figure 5B). Imaging of single-particles of hTR and mCherry-TRF1 telomeres was performed by constant acquisition at 100 msec/frame for a total of 10 sec. Single hTR particles were tracked using TrackMate, and colocalization with telomeres was quantified using CoPixie. With this acquisition mode, hTR particles colocalized with 48% of visible telomeres in cells overexpressing wild-type POT1, while the percentage of telomeres with hTR particles colocalization increased to 64% in cells expressing POT1-ΔOB (Figure 5C). Interestingly, a similar increase in hTR colocalization with telomeres was observed in cells expressing either POT1-Y223C (61% colocalization) or POT1-D224N (57% colocalization) (Figure 5C), showing that these mutants also increase telomerase access to telomeres.

**Figure 5:**
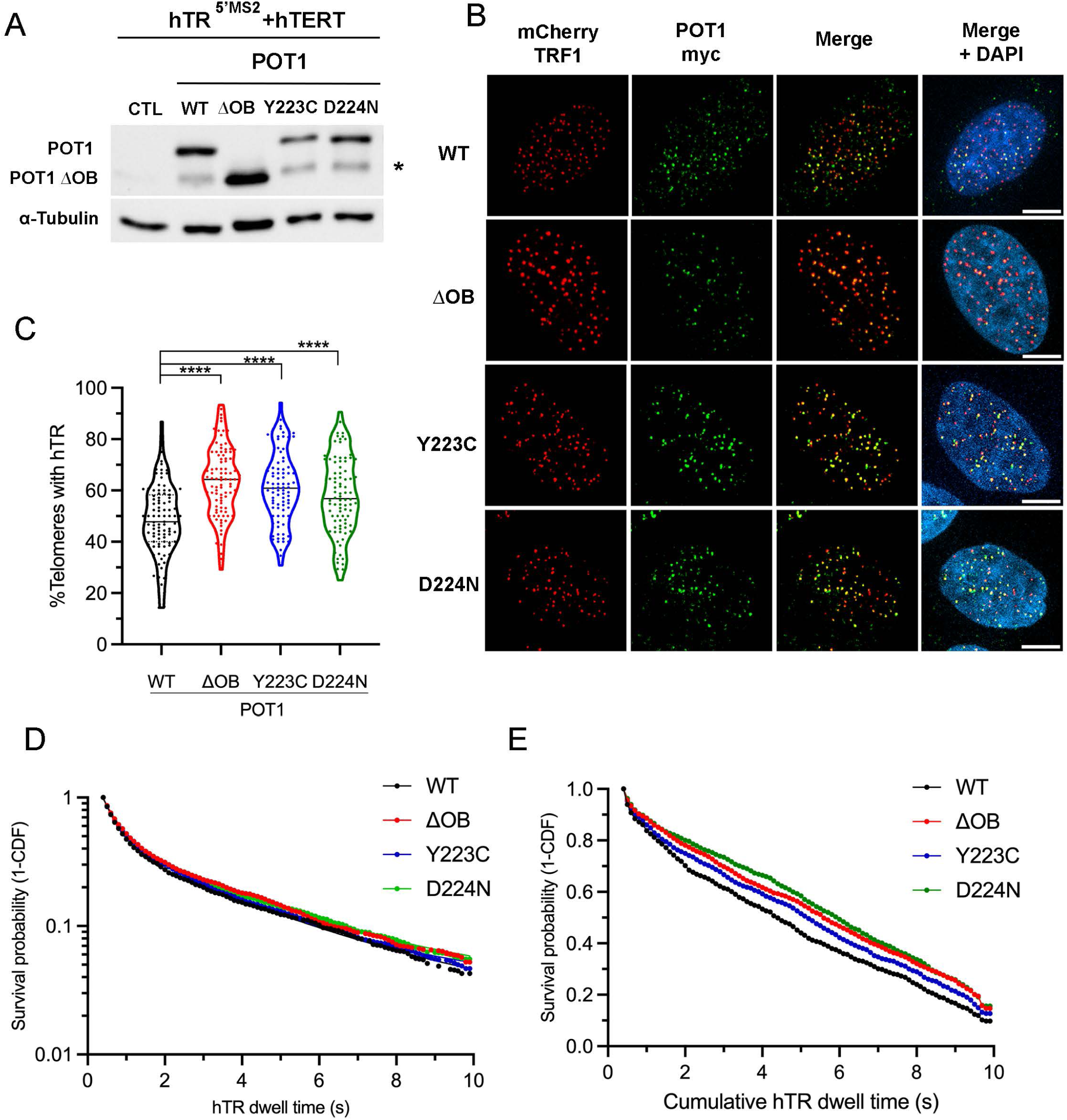
POT1 mutations Y223C and D224N increase telomerase access at telomeres. **A**) Western blot analysis to monitor expression level of myc-tagged POT1 wild-type (WT), POT1-ΔOB, POT1 Y223C and POT1-D224N in HeLa hTR^5’MS2^ cells. α-Tubulin was used as loading control. CTL: negative control. *: non-specific band. **B)** Immunofluorescence on myc-tagged POT1 WT and mutants in HeLa hTR^5’MS2^ cells. Immunostaining was performed with anti-myc antibody (green) and telomeres were detected with mCherry-TRF1 (red). DAPI: nuclear DNA. Scale bars: 8 μm. **C)** Percentage of telomeres colocalized with hTR in hTR^5’MS2^ + hTERT cells expressing POT1-WT, POT1-ΔOB, POT1 Y223C or POT1 D224N. ****p < 0.001. N = 85-95 cells. **D)** Survival probability analysis of individual hTR particles at telomeres in hTR^5’MS2^ + hTERT cells expressing POT1-WT, POT1-ΔOB, POT1 Y223C or POT1 D224N. Each curve represents data merged from >2700 tracks from 85-95 cells from 3 independent replicates. **E)** Survival probability analysis of the cumulative dwell-time of hTR particles at individual telomeres in hTR^5’MS2^ + hTERT cells expressing POT1-WT, POT1-ΔOB, POT1 Y223C or POT1 D224N. Each curve represents data merged from 1270 to 1460 tracks from 85-95 cells from 3 independent replicates. Statistical differences between groups were determined using Log-rank test: WT vs ΔOB:p<0.0001; WT vs Y223C: p=0.0112; WT vs D224N: p<0.0001.

The dwell-time of hTR particles at telomeres, and the cumulative dwell-time of all the hTR particles that visit a single telomere, were measured in each cell line. Fitting hTR dwell-time at telomeres in a survival probability plot using a two-phase decay equation revealed the residence time of hTR at telomeres in cells expressing POT1 WT or mutants. As shown in Figure 5D and Table S3, no significant difference in the dwell-time of hTR molecules at telomeres was observed between the various POT1 mutant and the wild-type. On the other hand, a significant increase in the measured cumulative dwell-time of hTR particles at single telomeres was observed between POT1 mutants and wild-type. Indeed, the median cumulative occupancy of telomeres increased from 4.4 sec in cells expressing POT1-WT to 5.6 sec in POT1-ΔOB, 5.2 sec with POT1-Y223C, and 6 sec with POT1-D224N (Figure 5E). Since the dwell-time of hTR molecules probing at telomeres is not changed in these POT1 mutants, these results suggest that the telomeric DNA is more accessible for hTR binding, increasing the cumulative dwell-time of hTR molecules at each telomere.

We switched to time-lapse acquisition with time intervals of 2 seconds between timepoints (0,5 Hz) to assess stable, long-lasting interactions between hTR particles and telomeres (See Supplementary Movies 5 to 6). CoPixie was used to quantify hTR-telomere colocalization events and dwell-time in cells expressing POT1-WT, POT1-ΔOB, or the mutants POT1-Y223C or POT1-D224N. This imaging mode confirmed the increased percentage of telomeres visited by hTR in cells expressing POT1-ΔOB, POT1-Y223C or POT1-D224N compared to cells expressing POT1-WT (Figure 6A). Interestingly, like POT1-K90E, POT1-Y223C increased the duration of long-lasting interactions between hTR and telomeres, from 32 seconds in POT1-WT to 47 seconds in POT-Y223C (Figure 6B and Table S4). On the other hand, POT1-D224N was more similar to POT1-ΔOB and POT1-WT, although long-lasting interactions above 150 sec displayed higher survival probabilities (Figure 6B and Table S4). Finally, cumulative hTR binding at telomeres revealed an increased median occupancy of telomeres by telomerase, from 28 sec in POT1-WT to 44 sec in POT1-ΔOB, 43 sec in POT1-Y223C and 36 sec in POT1-D224N (Figure 6C). Altogether, these results show that POT1-Y223C impacts both telomere accessibility and hTR long-lasting interactions at telomeres. Like POT1 K90E, this mutant differs from the POT1-ΔOB deletion, which affects only telomere accessibility.

**Figure 6:**
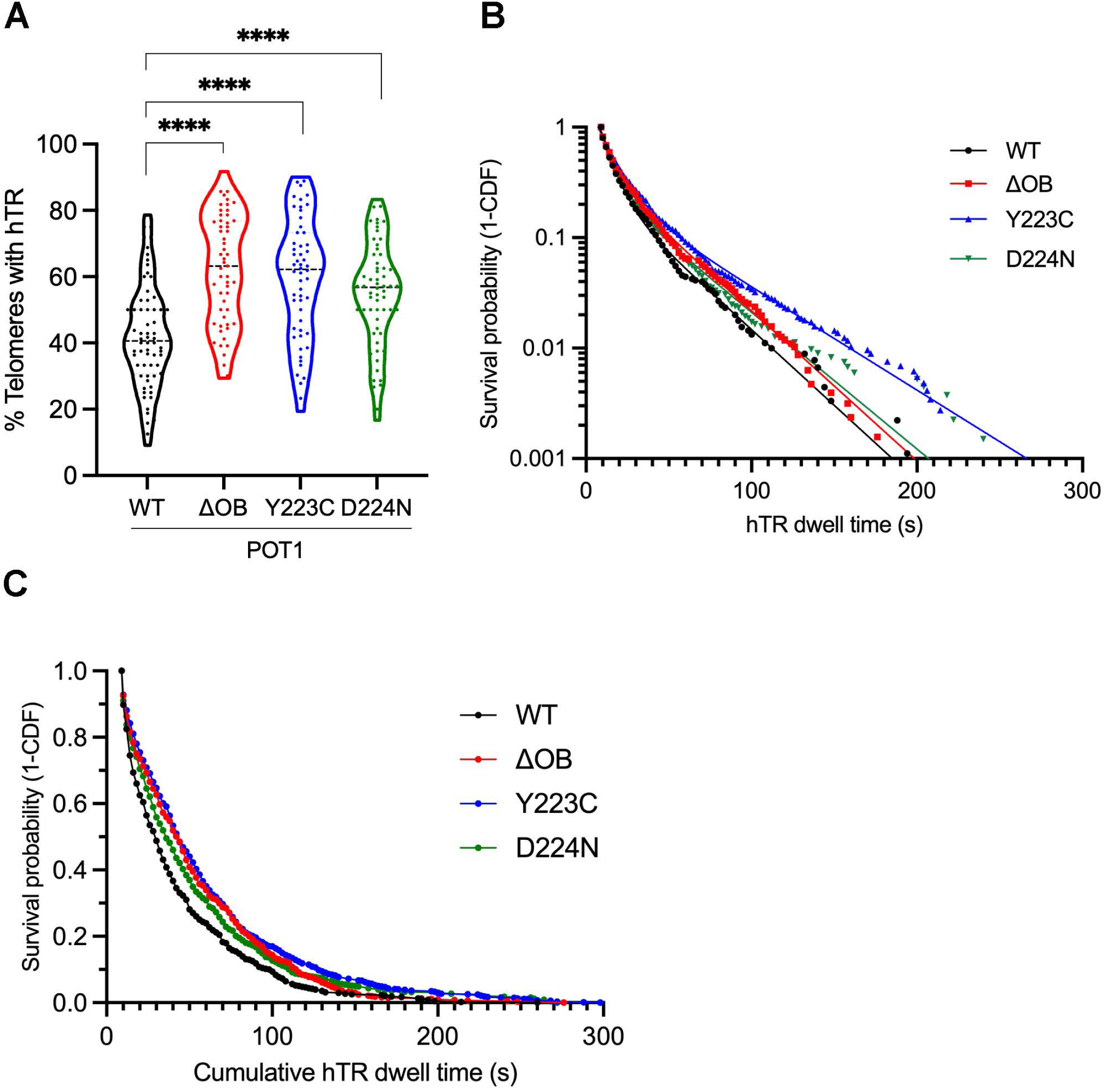
Long-term residence time of hTR at telomeres is increased in POT1 Y223C mutant. **A**) Percentage of telomeres colocalized with hTR in hTR^5’MS2^ + hTERT cells expressing POT1-WT, POT1-ΔOB, POT1 Y223C or POT1 D224N. ****p < 0.001. N = 55-66 cells. **B)** Survival probability analysis of individual hTR particles at telomeres in hTR^5’MS2^ + hTERT cells expressing POT1-WT, POT1-ΔOB, POT1 Y223C or POT1 D224N. Each curve represents data merged from 900 to 1300 tracks from 55 to 66 cells in 3 independent replicates. **C)** Survival probability analysis of the cumulative dwell-time of hTR particles at individual telomeres in hTR^5’MS2^ + hTERT cells expressing POT1-WT, POT1-ΔOB, POT1 Y223C or POT1 D224N. Each curve represents data merged from 500 to 700 tracks from 55 to 66 cells in 3 independent replicates. Statistical difference between groups was determined using Log-rank test: WT vs ΔOB: p<0.0001; WT vs Y223C: p<0.0001; WT vs D224N: p=0.0002.

## DISCUSSION

Quantification of colocalization events between single particles and their target could be time-consuming and limit the throughput of single-molecules colocalization analysis. In this study, we report the development of a new software called CoPixie, which can rapidly quantify colocalization events between a theoretically unlimited number of imaging channels, including single-particle movies. We show that CoPixie efficiently reproduced published results obtained by manual quantification of single-molecule imaging of telomerase interaction with telomeres (22). More importantly, CoPixie significantly decreases the time of analysis, with an average of 1.6 seconds per nucleus to complete an analysis of telomere occupancy and hTR dwell time at telomeres, versus 25 minutes per nucleus when analyzed manually.

We applied multicolor single-molecule imaging and analysis with CoPixie to study the role of POT1 in regulating telomerase-telomeres interactions and dynamics. We used different modes of acquisition to measure the dynamics of telomerase-telomeres interactions. Rapid acquisition at 10 Hz allows the detection of shorter or probing telomerase-telomeres interactions mediated mainly by hTERT-TPP1 interactions (20). Acquisition at 0.5 Hz allows the capture of long-lasting, or engaged interactions mediated by hTR binding to telomeric ssDNA (22). Movies acquired via both modes of acquisition were processed with CoPixie to detect and quantify hTR-telomere interactions and dwell-time. With both modes of acquisition, CoPixie reproducibly reported an increase in 1) hTR-telomere associations and 2) cumulative dwell-time of hTR at telomeres, in cells expressing POT1-ΔOB compared to cells expressing POT1-WT, as previously published (22). Finally, we also tested two independent cell lines generated by infection of HeLa hTR^5’MS2^ cells with lentivirus expressing POT1-WT or POT1-ΔOB (Figures 3A and 5A). Imaging of the two pairs of cell lines at separate times, and analysis by CoPixie, reproducibly quantified a significant increase in hTR-telomere colocalizations in POT1-ΔOB cells compared to POT1-WT cells (Figures 3B and 5B). Altogether, these results show that CoPixie is a robust and reproducible pipeline for single-particle track colocalization analyses.

We further applied our imaging analysis pipeline to study the impact of overexpression of POT1 and cancer-associated POT1 mutations on telomerase-telomeres interaction and dynamics. First, we observed that overexpression of POT1 itself increased the dwell-time of short-lived, or probing, interactions between telomerase and telomeres. These probing events are possibly mediated by hTERT-TPP1 interactions since the deletion of POT1 N-terminal OB-fold domain had no additional impact on these binding events. Since POT1 is a limiting member of the Shelterin complex at telomeres (28), and can directly interacts with hTERT while in complex with TPP1 (37), increasing POT1 association with TPP1 at telomeres by overexpression of POT1 may help stabilize telomerase probing interactions. The longer dwell-time of telomerase probing events, which also result in longer cumulative dwell-time of telomerase at telomeres, could increase the probability that telomerase becomes engaged in elongation events. This may explain why overexpression of POT1 can lead to the elongation of telomeres, as previously reported in several articles (35,38–40).

Secondly, we found that expression of the cancer-associated mutations POT1-K90E, Y223C and D224N, increases the recruitment of telomerase at telomeres. Indeed, the percentage of telomeres with colocalized hTR, and the cumulative dwell-time of hTR molecules at telomeres, increased in the K90E, Y223C and D224N mutants. Remarquably, even the K90E mutation, which does not affect the capacity of POT1 to bind telomeric ssDNA *in vitro* (35), increases the accessibility of telomeres for telomerase binding *in vivo*. Although the mechanism remains unclear, we can speculate that the POT1 K90E mutant may affect other layers of regulation of telomerase binding at telomeres, such as ATR or ATM signalling, for instance (22,35,41). Unexpectedly, unlike POT1-ΔOB, the K90E and Y223C mutations also increase the residence time of the long-lasting hTR particles at telomeres. Since these long-lasting interactions depend on the base-pairing of hTR to the 3’overhang of telomeres, leading to the elongation of telomeres (21,22), this suggests that these mutations may impact an activity that limits telomerase elongation. Such an activity may lie in the CST complex, which acts as a terminator of telomerase elongation at telomeres (42) and is known to interact with POT1 (42,43). Interestingly, the K90E mutant was reported to disrupt the interaction of the CST complex with telomeres (35), which could lead to a reduction of the inhibitory activity of CST on elongating telomerase. This could explain why long-lasting interactions between telomerase and telomeres are particularly affected in the K90E mutant. Regarding the POT1 Y223C mutant, it remains unclear how it could affect long-lasting interactions of telomerase with telomeres. In future work, it will be interesting to determine if, like the K90E mutant, the POT1-Y223C also impacts the CST complex at telomeres.

CoPixie helps solve some of the limitations of current approaches based on single-particle tracking, which rely on changes in diffusion coefficient and residence time to infer binding. It combines particle tracks and object-based colocalization to accurately detect the start and end of interactions between particles *in vivo*. Copixie is remarkably flexible in the types of inputs it accepts, because single images can be used, as well as user drawn regions. Another strength of the CoPixie pixel-based colocalization algorithm is that it is not limited to pairs of objects, *i.e*. two hTR particles can be detected simultaneously on the same telomere. CoPixie colocalisation analysis is only limited by the accuracy of single-particle tracking in each channel. We think that CoPixie can be used to explore a variety of questions involving macromolecular interactions in live-cell imaging, from multi-wavelength single-particle tracks, to the quantification of particles within specific cellular subdomains.

## DATA AVAILABILITY

CoPixie can be installed from https://github.com/drs/CoPixie. All data are available upon request.

## SUPPLEMENTARY DATA

Supplementary Data are available.

## ACKNOWLEDGEMENTS

We thank Agnel Sfeir for lentiviral vectors for expression of myc-tagged POT1. *Authors contributions*: S.P, K.M and E.Q developed CoPixie; S.P wrote CoPixie; K.M, M.F, A.D and E.Q performed experiments; K.M, M.F, E.Q and P.C analyzed the data; and S.P, E.Q and P.C wrote the manuscript.

## Supplementary Tables

**Table S1:** Residence time (Tau) and half-life of fast and slow hTR particles at telomeres in cells expressing POT1-WT, ΔOB or K90E mutants, or empty vector. Data are from 10 seconds acquisitions at 10 Hz. Mean ±95%CI are reported. ****p<0.0001: empty versus WT (related to Figure 3D).

| Cell line | Nuclei count | Tau (fast) | Tau (slow) | Half life (fast) | Half life (slow) |
| --- | --- | --- | --- | --- | --- |
| Empty | 61 | 0,29 ± 0,01**** | 1,95 ± 0,11**** | 0,21 ± 0,01**** | 1,35 ± 0,08**** |
| WT | 64 | 0,44 ± 0,02 | 2,75 ± 0,27 | 0,30 ± 0,017 | 1,91 ± 0,19 |
| ΔOB | 65 | 0,39 ± 0,01 | 2,91 ± 0,14 | 0,27 ± 0,01 | 2,02 ± 0,1 |
| K90E | 58 | 0,38 ± 0,008 | 2,97 ± 0,09 | 0,26 ± 0,006 | 2,06 ± 0,06 |

**Table S2:** Residence time (Tau) and half-life of fast and slow hTR particles at telomeres in cells expressing POT1-WT, ΔOB or K90E mutants, or empty vector. Data are from 300 seconds acquisitions at 0.5 Hz. Mean ±95%CI are reported. ****p<0.0001: K90E versus WT (related to Figure 4B).

| Cell line | Nuclei count | Tau (fast) | Tau (slow) | Half life (fast) | Half life (slow) |
| --- | --- | --- | --- | --- | --- |
| Empty | 55 | 7,16 ± 0,5 | 38,47 ± 1,7 | 4,96 ± 0,36 | 26,66 ± 1,2 |
| WT | 62 | 7,6 ± 0,9 | 38,14 ± 1,8 | 5,27 ± 0,6 | 26,44 ± 1,2 |
| ΔOB | 52 | 5,86 ± 1,7 | 30,95 ± 2,1 | 4,06 ± 1,2 | 21,45 ± 1,5 |
| K90E | 54 | 10,06 ± 0,67**** | 51,57 ± 1,8**** | 6,98 ± 0,46**** | 35,75 ± 1,2**** |

**Table S3:** Residence time (Tau) and half-life of fast and slow hTR particles at telomeres in cells expressing POT1-WT, ΔOB, Y223C or D224N mutants. Data are from 10 seconds acquisitions at 10 Hz. Mean ±95%CI are reported. (related to Figure 5D).

**(related to Figure 5D).**
| Cell line | Nuclei count | Tau (fast) | Tau (slow) | Half life (fast) | Half life (slow) |
| --- | --- | --- | --- | --- | --- |
| WT | 95 | 0,398±0,016 | 3,39 ± 0,22 | 0,28 ± 0,011 | 2,35 ± 0,15 |
| ΔOB | 95 | 0,428±0,015 | 3,53 ± 0,16 | 0,30 ± 0,01 | 2,45 ± 0,11 |
| Y223C | 85 | 0,394±0,02 | 2,91 ± 0,15 | 0,27 ± 0,014 | 2,01 ± 0,1 |
| D224N | 90 | 0,405±0,026 | 3,33 ± 0,27 | 0,28 ± 0,018 | 2,31 ± 0,19 |

**Table S4:** Residence time (Tau) and half-life of fast and slow hTR particles at telomeres in cells expressing POT1-WT, ΔOB, Y223C or D224N mutants. Data are from 300 seconds acquisitions at 0.5 Hz. Mean ±95%CI are reported. ****p<0.0001: Y223C versus WT (related to Figure 6B).

| Cell line | Nuclei count | Tau (fast) | Tau (slow) | Half life (fast) | Half life (slow) |
| --- | --- | --- | --- | --- | --- |
| WT | 66 | 7,57 ± 1,17 | 31,65 ± 3,4 | 5,25 ± 0,81 | 21,94 ± 2,36 |
| ΔOB | 55 | 7,27 ± 1,08 | 31,96 ± 1,71 | 5,04 ± 0,75 | 22,15 ± 1,18 |
| Y223C | 56 | 11,19 ± 1,2**** | 46,52 ± 3,24**** | 7,76 ± 0,83**** | 32,25 ± 2,25**** |
| D224N | 60 | 7,927 | 35,08 | 5,494 | 24,32 |

